# Chronic environmental or genetic elevation of galanin in noradrenergic neurons confers stress resilience in mice

**DOI:** 10.1101/2020.04.24.060608

**Authors:** Rachel P. Tillage, Genevieve E. Wilson, L. Cameron Liles, Philip V. Holmes, David Weinshenker

**Affiliations:** Department of Human Genetics, Emory University School of Medicine, Atlanta, GA 30322, USA; Department of Psychology, University of Georgia, Athens, GA 30602, USA

## Abstract

The neuropeptide galanin has been implicated in stress-related neuropsychiatric disorders in both humans and rodent models. While pharmacological treatments for these disorders are ineffective for many individuals, physical activity is beneficial for stress-related symptoms. Galanin is highly expressed in the noradrenergic system, particularly the locus coeruleus (LC), which is dysregulated in stress-related disorders and activated by exercise. Galanin expression is elevated in the LC by chronic exercise, and blockade of galanin transmission attenuates exercise-induced stress resilience. However, most research on this topic has been done in rats, so it is unclear whether the relationship between exercise and galanin is species-specific. Moreover, use of intracerebroventricular galanin receptor antagonists in prior studies precluded defining a causal role for LC-derived galanin specifically. Therefore, the goals of this study were twofold. First, we investigated whether physical activity (chronic voluntary wheel running) increases stress resilience and galanin expression in the LC of mice. Next, we used transgenic mice that overexpress galanin in noradrenergic neurons (Gal OX) to determine how chronically elevated noradrenergic-derived galanin, alone, alters anxiogenic-like responses to stress. We found that three weeks of *ad libitum* access to a running wheel in their home cage increased galanin mRNA in the LC of mice and conferred resilience to a stressor. The effects of exercise were phenocopied by galanin overexpression in noradrenergic neurons, and Gal OX mice were resistant to the anxiogenic effect of optogenetic LC activation. Together, these findings support a role for chronically increased noradrenergic galanin in mediating resilience to stress.

**Significance statement:** Understanding the neurobiological mechanisms underlying behavioral responses to stress is necessary to improve treatments for stress-related neuropsychiatric disorders. Increased physical activity is associated with stress resilience in humans, but the neurobiological mechanisms underlying this effect are not clear. Here we investigate the anxiolytic potential of the neuropeptide galanin from the main noradrenergic nucleus, the locus coeruleus (LC). We show that chronic voluntary wheel running in mice galanin expression in the LC and stress resilience. Furthermore, we show that genetic overexpression of galanin in noradrenergic neurons confers resilience to the anxiogenic effects of foot shock and optogenetic LC activation. These findings support a role for chronically increased noradrenergic galanin in mediating resilience to stress.

## INTRODUCTION

Stress-related neuropsychiatric disorders affect approximately 600 million people worldwide, yet current pharmacological treatments have limited efficacy and cause adverse side effects for many people (Nestler et al., 2002; Rush et al., 2006; James et al., 2018). Clinical studies have consistently linked physical exercise to improvements in a wide array of neuropsychiatric disorders (Herring et al., 2010; Cooney et al., 2013; Ashdown-Franks et al., 2020). Individuals who regularly exercise are less likely to experience stress-related neuropsychiatric disorders, such as depression, anxiety, and post-traumatic stress disorder (Whitworth and Ciccolo, 2016; Chekroud et al., 2018; Harvey et al., 2018), and chronic voluntary wheel running increases resilience to various stressors in rodents (Sciolino et al., 2012; Kingston et al., 2018; Mul et al., 2018; Tanner et al., 2019). A multitude of biological changes occur as a result of chronically increased physical activity which may or may not be causally linked to alterations in stress resilience and mood; however, a promising candidate for mediating these beneficial effects is the neuropeptide galanin.

Galanin is abundant in the brain (Tatemoto, 1983; Kofler et al., 2004; Fang et al., 2015), and modulates stress, mood, cognition, food intake, nociception, and seizures (Mitsukawa et al., 2008; Lang et al., 2015). Human studies have implicated genetic variants in the genes encoding the galanin gene and its receptors (GalR1, 2, and 3) in conferring an increased risk of depression and anxiety, especially in the context of environmental stress exposure, and postmortem studies have similarly revealed galaninergic dysregulation in people with major depressive disorder (Barde et al., 2016).

While the pattern of galanin expression in the brain can vary from species to species, it is particularly rich in the noradrenergic locus coeruleus (LC) of both humans and rodents (Skofitsch, 1985; Melander, 1986; Holets et al., 1988; Chan-Palay et al., 1990; Perez et al., 2001; Le Maitre et al., 2013). The LC is activated by stress, and norepinephrine (NE) release plays a crucial role in the stress response, coordinating a host of downstream effects via broad axonal projections required for the “fight-or-flight” response (Valentino and Van Bockstaele, 2008). Consistent with this role, the LC-NE system is dysregulated in stress-related neuropsychiatric disorders, such as anxiety, depression, and post-traumatic stress disorder (Roy et al., 1988; Ordway et al., 1994; Wong et al., 2000; Bissette et al., 2003; Ehnvall et al., 2003; Roy et al., 2017; Naegeli et al., 2018). In rats, galanin expression increases in the LC following chronic wheel running and correlates with running distance (Van Hoomissen et al., 2004; Holmes et al., 2006; Sciolino et al., 2012; Epps et al., 2013; Sciolino et al., 2015), and in Sciolino et al. (2015), we found that wheel running confers resilience to a stressor. Galanin has both acute neuromodulatory and chronic neurotrophic properties, and the beneficial effects of exercise in rats were blocked by chronic, but not acute, intracerebroventricular (ICV) infusion of a galanin antagonist and recapitulated by chronic ICV infusion of galanin alone, suggesting a neurotrophic mechanism of action (Sciolino et al., 2015). However, it is not known whether these phenomena can be extrapolated to other species or attributed to LC-derived galanin.

To investigate the contribution of chronically elevated noradrenergic galanin to stress resilience in mice, we investigated behavioral responses to foot shock in both an environmental (voluntary wheel running) and genetic (dopamine β-hydroxylase-galanin transgene; “Gal OX”) model of LC galanin overexpression. To isolate anxiety-like responses mediated specifically by the LC, we also examined the behavior of Gal OX mice following optogenetic LC activation. Our findings support a causal role for chronically elevated noradrenergic-derived galanin in promoting stress resilience.

## MATERIALS AND METHODS

### Animals

All procedures related to the use of animals were approved by the Institutional Animal Care and Use Committee of Emory University and were in accordance with the National Institutes of Health guidelines for the care and use of laboratory animals. All mice were group housed and maintained on a 12/12-h light-dark cycle with access to food and water *ad libitum*, unless noted otherwise. All manipulations and behavioral tests occurred during the light cycle. Adult male and female mice (3-9 months), with equal numbers of both sexes, were used for all experiments. The Gal OX mice were generated with a transgene containing the mouse galanin gene driven by the human *dopamine β-hydroxylase* (*Dbh*) promoter, resulting in an approximately 5-fold increase in galanin mRNA in noradrenergic and adrenergic neurons and a 2-fold increase in galanin protein in LC-innervated forebrain regions compared to wild-type (WT) littermates (Jackson Labs stock #004996) (Steiner et al., 2001). All mice used in this study were on a C57BL/6J background.

### Exercise

Both exercise and sedentary mice were singly housed for one week prior to the start of running wheel experiments. On the first day of the experiment, low-profile wireless running wheels (Med Associates, St. Albans, VT) were placed in the cage of the exercise mice and monitored for 3 weeks. All mice were weighed once a week.

### Behavioral assays

Baseline behavioral assays (open field (OF), elevated plus maze (EPM), forced swim test (FST), fear conditioning, nestlet shredding, elevated zero maze (EZM), marble burying, novelty-suppressed feeding (NSF), locomotor activity, and shock-probe defensive burying (SPDB)) were conducted as previously described (Tillage et al., 2020) with at least 4-5 d between tests. For the behavioral battery following exercise, the assays were conducted 24-48 h post-foot shock stress, with at least 2-3 h between tests, in order of least stressful to most stressful to minimize effects from the previous tests.

### Stress paradigm

The foot shock stress protocol was modified from a previously published paradigm (Lecca et al., 2016). Mice were individually exposed to 20 min of foot shock exposure consisting of 19 shocks randomly interspaced by 30, 60, or 90 s (0.5-ms shocks, 1 mA) in chambers (Coulbourn Instruments, Holliston, MA) equipped with a house light, a ceiling-mounted camera, and an electric grid shock floor. Control animals were placed in the chamber for 20 min but were not administered shocks. Chambers were cleaned with MB-10 between animals.

### Corticosterone (CORT) measurement

Mice used for CORT measurement went through the foot shock stress paradigm described above and were anesthetized with isoflurane 15 min after the end of the stress. Mice were rapidly decapitated, and trunk blood was collected in EDTA-coated tubes (Sarstedt Inc., Newton, NC) and chilled on ice. Blood was centrifuged for 20 min at 3000 rpm at 4°C, and the resulting plasma was collected and stored at -80°C. CORT was measured using the Enzo Life Sciences kit (Farmingdale, NY) following the manufacturer’s small volume protocol for blood plasma, including diluting samples 1:40 with a steroid displacement reagent solution.

### Galanin *in situ* hybridization and densitometry

To measure galanin mRNA in the LC following exercise, mice were deeply anesthetized with isoflurane and rapidly decapitated for brain extraction. Brains were collected, flash frozen in Tissue Freezing Medium, and stored at -80°C until processing. Brains were sectioned at 12-µm and collected on gelatin-coated slides (SouthernBiotech, Birmingham, AL). Slides were stored at -80°C. Galanin *in situ* hybridization was conducted and images were acquired as previously described (Sciolino et al., 2015). For quantification, images were converted to 16-bit in NIH ImageJ (https://imagej.nih.gov/ij/) and the mean grayscale value was measured in 3-4 LC sections per animal using a standardized region of interest to obtain an average measurement per animal. The mean grayscale value of the background was subtracted out for each measurement. The experimenter was blind to treatment during image analysis.

### Stereotaxic surgery

For optogenetic experiments, mice were anesthetized with isoflurane and given the analgesic meloxicam at the start of the surgery (5 mg/kg, s.c.). A lentiviral vector containing the channelrhodopsin (ChR2) construct with an mCherry tag under control of the noradrenergic-specific PRSx8 promoter (Hwang et al., 2001) was infused unilaterally into the LC (−5.4AP; +1.2ML; -4.0DV) with a 5 µL Hamilton syringe and Quintessential Stereotaxic Injector (Stoelting, Wood Dale, IL) pump at a rate of 0.15 µL/min. Unilateral LC stimulation is sufficient to produce behavioral effects in mice that are indistinguishable from bilateral stimulation (Carter et al., 2010; McCall et al., 2015). Control mice received a lentivirus containing mCherry alone under the PRSx8 promoter. Each animal received a 0.7 µL infusion and the infusion needle was left in place for 5 min after infusion to allow for viral diffusion. An optic fiber ferrule (ThorLabs, Newton, NJ) was implanted 0.5 mm dorsal to the viral injection site (−5.4AP; +1.2ML; -3.5DV) and permanently attached to the skull with screws and dental acrylic. Mice were singly housed after this surgery to prevent cage mates from damaging the headcap and given at least 3 weeks to recover and allow for full viral expression before testing.

### Optogenetic stimulation

Mice were habituated to handling and connection of the optic patch cable to the implanted optic ferrule for at least one week prior to testing. The LC optogenetic stimulation was based on a previously published paradigm (McCall et al., 2015). Photostimulation was delivered to the LC for 30 min (5 Hz tonic stimulation, 10 ms light pulses, 473 nm) in 3 min on/off bins in the home cage.

### Histology

To assess correct targeting of the LC in mice used for optogenetic experiments, all animals were exposed to 5 Hz LC photostimulation for 15 min in the home cage, then anesthetized 90 min later with isoflurane and transcardially perfused with potassium phosphate-buffered saline (KPBS), followed by 4% PFA in PBS. Brains were postfixed overnight by immersion in 4% PFA at 4°C, and then transferred to 30% sucrose in KPBS for 48 h at 4°C. Brains were flash frozen in isopentane on dry ice and embedded in Tissue Freezing Medium. Tissue was cryosectioned at 40-µm for immunohistochemistry. Viral expression and correct optic fiber targeting were assessed by immunostaining for the mCherry tag using rabbit anti-DsRed primary antibody (1:1000) and goat anti-rabbit 568 secondary (1:500). Sections were co-stained for the noradrenergic marker tyrosine hydroxylase (TH) with chicken anti-TH (1:1000) and goat anti-chicken 488 secondary (1:500). In adjacent LC sections, activated LC neurons were detected with rabbit anti c-Fos primary antibody (1:5,000) with goat anti-rabbit 488 secondary (1:500) and TH-expressing cells were co-stained using chicken anti-TH (1:1000) with goat anti-chicken 568 secondary (1:500). After staining, all sections were mounted on slides and cover slipped with Fluoromount plus DAPI (Southern Biotech). Images were collected on a Leica DM6000B epifluorescent upright microscope at 10x or 20x. Mice were excluded from analyses if they did not show viral expression in the LC, correct optic fiber targeting, and increased c-Fos expression in the LC compared to the control animals of the same cohort. Antibodies are summarized in Table 1.

**Table 1.** Antibodies used.

| **Antibody** | **Species** | **Dilution** | **Source** | **Catalog #** |
| --- | --- | --- | --- | --- |
| TH | Chicken | 1:1000 | Abcam | AB76442 |
| c-fos | Rabbit | 1:5000 | Millipore | ABE457 |
| DsRed | Rabbit | 1:1000 | Takara | 632496 |
| Alexa Fluor 488 anti-rabbit | Goat | 1:500 | Invitrogen | A-11008 |
| Alexa Fluor 568 anti-rabbit | Goat | 1:500 | Invitrogen | A-11011 |
| Alexa Fluor 488 anti-chicken | Goat | 1:500 | Abcam | AB150169 |
| Alexa Fluor 568 anti-chicken | Goat | 1:500 | Invitrogen | A-11041 |

### Statistical analysis

Data were analyzed via unpaired t-test or two-way ANOVA with Sidak’s correction for multiple comparisons, where appropriate. Significance was set at p<0.05 and two-tailed variants of tests were used throughout. Data are presented as mean ± SEM. Calculations were performed and figures created using Prism Version 8 (GraphPad Software, San Diego, CA).

## RESULTS

### Wheel running characteristics

Singly housed WT C57BL6/J mice were given unrestricted access to a running wheel in their home cage for 21 d. Mice steadily increased their running distance over the first week, after which they ran approximately 10-16 km per day, which is comparable to distances seen for C57BL/6J mice in previous studies using similar running wheels (De Bono et al., 2006; Goh and Ladiges, 2015) (**Fig. 1a**). Female mice ran significantly more than males (F_1,12_=9.101, p=0.0107). Male exercise mice lost weight over the course of the 3 weeks compared to their sedentary counterparts, which tended to gain weight over time (F_1,12_=8.441, p=0.0132) (**Fig. 1b**). Female exercise mice did not show weight loss, but did have a significant slowing of normal weight gain compared to the sedentary females (F_1,13_=13.69, p=0.0027) (**Fig. 1c**).

**Figure 1.**
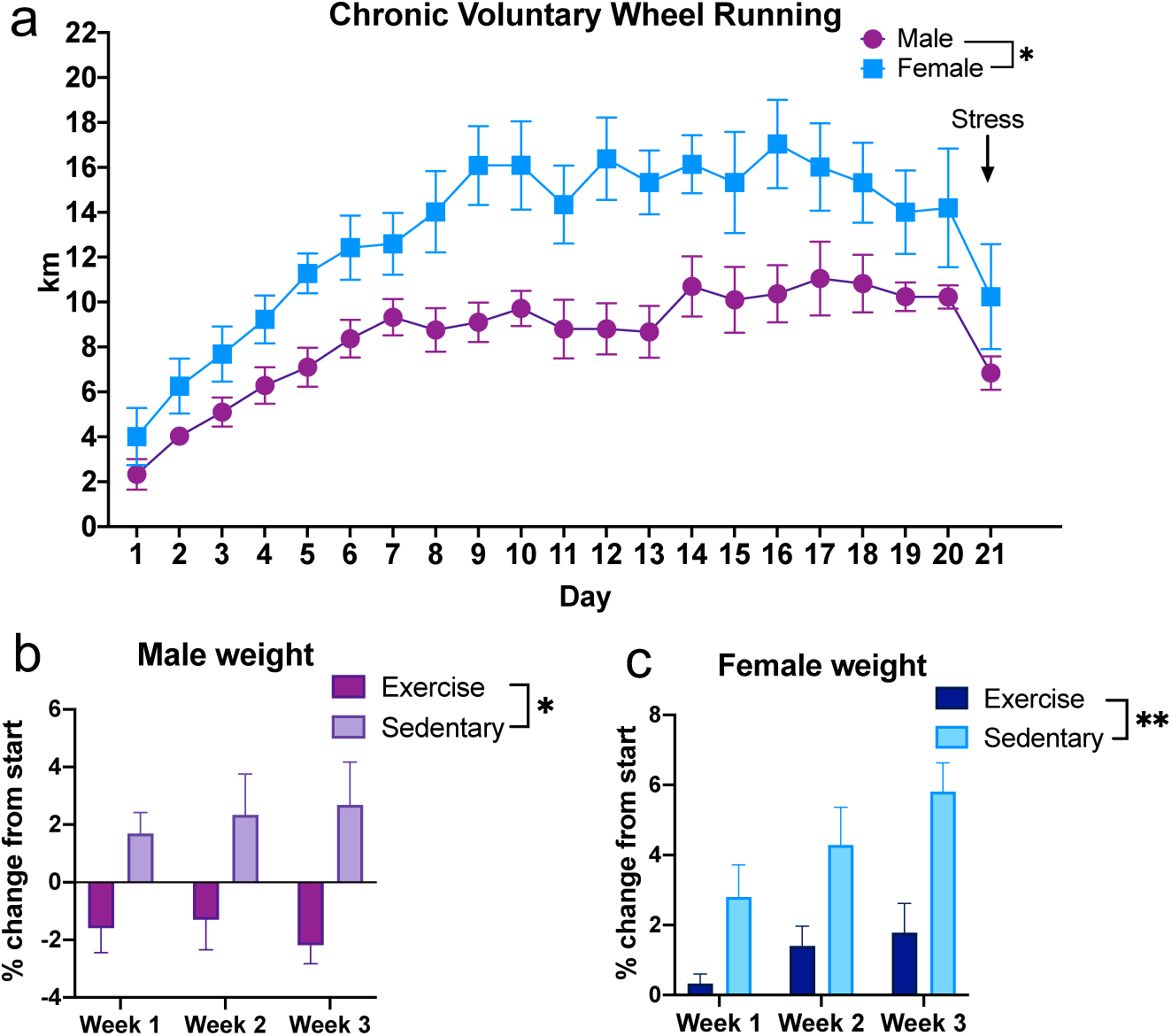
Distance traveled and weight change in exercising mice. Wild-type C57Bl6/J mice were given *ad libitum* access to running wheels in their home cage for 3 weeks. (a) Female mice consistently ran farther distances per day than male mice over three weeks. (b) Male exercise lost weight over time compared to their sedentary counterparts. (c) Female exercise mice gained weight more slowly than their sedentary counterparts. *n* = 7-8 mice per group. Error bars show SEM. *p<0.05, **p<0.01.

### Exercise increases resilience to foot shock stress

At the end of the 3 weeks, half of the mice were subjected to a foot shock stressor. The following day, mice were put through a battery of behavioral tasks consisting of elevated zero maze (EZM), marble burying (MB), and shock-probe defensive burying (SPDB). Mice were then food-deprived overnight and tested in a novelty-suppressed feeding (NSF) assay the next day before euthanasia and tissue collection (**Fig. 2a**). For the EZM, a two-way ANOVA showed a significant exercise x stress interaction (F_1,25_=5.718, p=0.0246), and post hoc tests revealed that sedentary mice had a significant decrease in the time spent in the open arms of the EZM after stress (p=0.0294), while stress had no effect on the exercise animals (p=0.6810) (**Fig. 2b**). No differences were seen in the MB test as a result of exercise (F_1,25_=2.265, p=0.1448) or stress (F_1,25_=0.06874, p=0.7953) (**Fig. 2c**).

**Figure 2.**
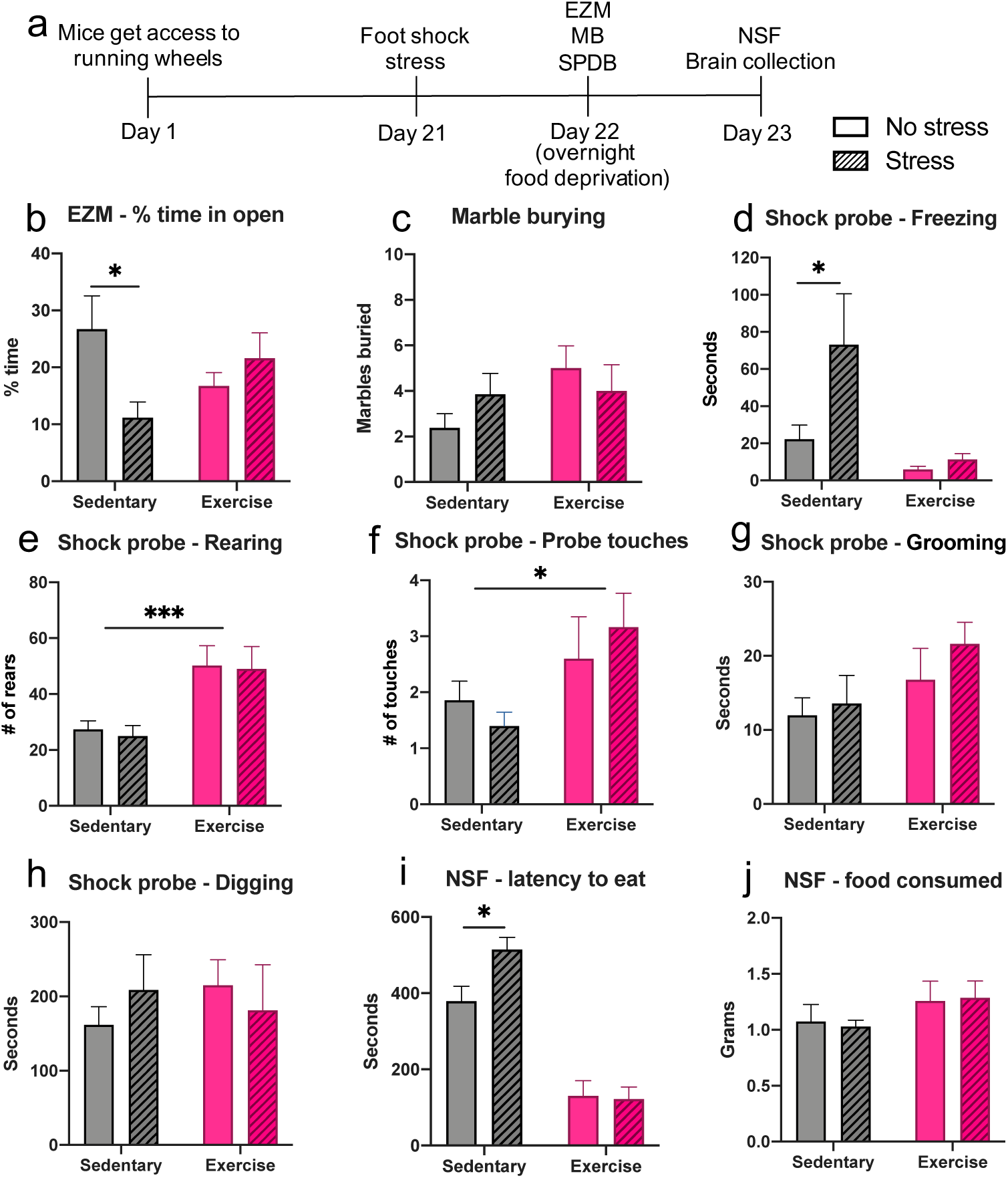
Exercise increases resilience to foot shock stress. Wild-type C57Bl6/J mice were given *ad libitum* access to running wheels in their home cage (“Exercise”) or no running wheel (“Sedentary”) for 3 weeks, then were tested in behavioral assays 24-48 h following foot shock (“Stress”) or no foot shock (“No Stress”). (a) Exercise paradigm timeline. EZM, elevated zero maze. MB, marble burying. SPDB, shock probe defensive burying. NSF, novelty-suppressed feeding. (b) Sedentary mice show decreased time spent in the open arms of the EZM after stress, but exercise mice do not. (c) No significant differences were seen in the MB assay. (d) In the SPDB assay, sedentary stressed mice showed increased freezing compared to sedentary non-stressed mice, but there was no difference between stressed and non-stress exercise mice. (e) Exercise mice showed an overall increase in both rearing and (f) number of probe touches compared to sedentary mice, with no effect of stress. (g) There were no differences as a result of exercise or stress on grooming or (h) digging in the SPDB assay. (i) In the NSF test, sedentary mice took significantly longer to eat after stress compared to non-stressed sedentary mice, with no difference between stressed and non-stressed exercise mice. (j) Exercise mice tended to consume more food during the hour after the NSF test, but it was not significant. *n* = 7-8 mice per group. Error bars show SEM. *p<0.05, ***p<0.001.

In the SPDB assay, a two-way ANOVA showed a main effect of exercise (F_1,19_=9.133, p=0.007) and stress (F_1,19_=4.759, p=0.0419) on freezing behavior, with the stressed sedentary animals spending significantly more time freezing compared to non-stressed sedentary animals (p=0.021). The effects of stress were abrogated by exercise (p=0.9489) (**Fig. 2d**). There was a main effect of exercise on rearing behavior, showing that regardless of stress exposure (F_1,19_=0.09776, p=0.7579), exercise mice displayed more rearing bouts in the SPDB task (F_1,19_=16.24, p=0.0007) (**Fig. 2e**). Exercise mice also had a slight, but significant increase in the number of shock probe touches from the electrified probe compared to sedentary mice (F_1,19_=6.004, p=0.0241), regardless of stress exposure (F_1,19_=0.01144, p=0.9160) (**Fig. 2f**).

There was no effect of exercise or stress on the amount of time mice spent grooming (exercise: F_1,19_=3.917, p=0.0625; stress: F_1,19_=0.9914, p=0.3319) or digging (exercise: F_1,19_=0.08843, p=0.7694; stress: F_1,19_=0.02293, p=0.8812) in the SPDB test (**Fig. 2g-h**).

In the NSF task, there was a main effect of exercise on the latency to eat the food in the novel environment (F_1,25_=78.94, p<0.0001) but no main effect of stress (F_1,25_=3.092, p=0.0909). Post hoc testing revealed that stressed sedentary animals had a significantly longer latency to eat compared to non-stressed sedentary animals (p=0.0243), with no effect of stress on the exercise animals (p=0.9825) (**Fig. 2i**). As a control, the amount of food consumed by each mouse in 1 h immediately after the NSF test was recorded. There were no significant differences due to exercise (F_1,25_=2.371, p=0.1361) or stress (F_1,25_=0.003919, p=0.9506) (**Fig. 2j**). Together, these results demonstrate that chronic wheel running affords protection from the anxiogenic-like effects of foot shock stress.

### Exercise increases gal mRNA in the LC

Several previous studies have shown that chronic voluntary exercise increases prepro-galanin mRNA in the LC of rats (Van Hoomissen et al., 2004; Holmes et al., 2006; Sciolino et al., 2012; Sciolino et al., 2015), but this change has never been examined in mice. We measured prepro-galanin mRNA in the LC of exercise and sedentary mice using *in situ* hybridization and found a robust and significant increase in galanin mRNA in exercise mice compared to their sedentary counterparts (F_1,23_=58.24, p<0.0001), with no difference according to stress exposure (F_1,23_=0.08014, p=0.7796) (**Fig. 3a-b**). Furthermore, we found a positive correlation between the average distance each mouse ran per day in the third week and the level of galanin mRNA in the LC (r^2^=0.4196, p=0.0312) (**Fig. 3c**), but not in week 1 (r^2^=0.08222, p=0.3926) or week 2 (r^2^=0.2234, p=0.142) (**Fig. 3d-e**). These results indicate that elevation of galanin expression in the LC occurs as a result of chronic physical activity in mice, similar to what is observed in rats.

**Figure 3.**
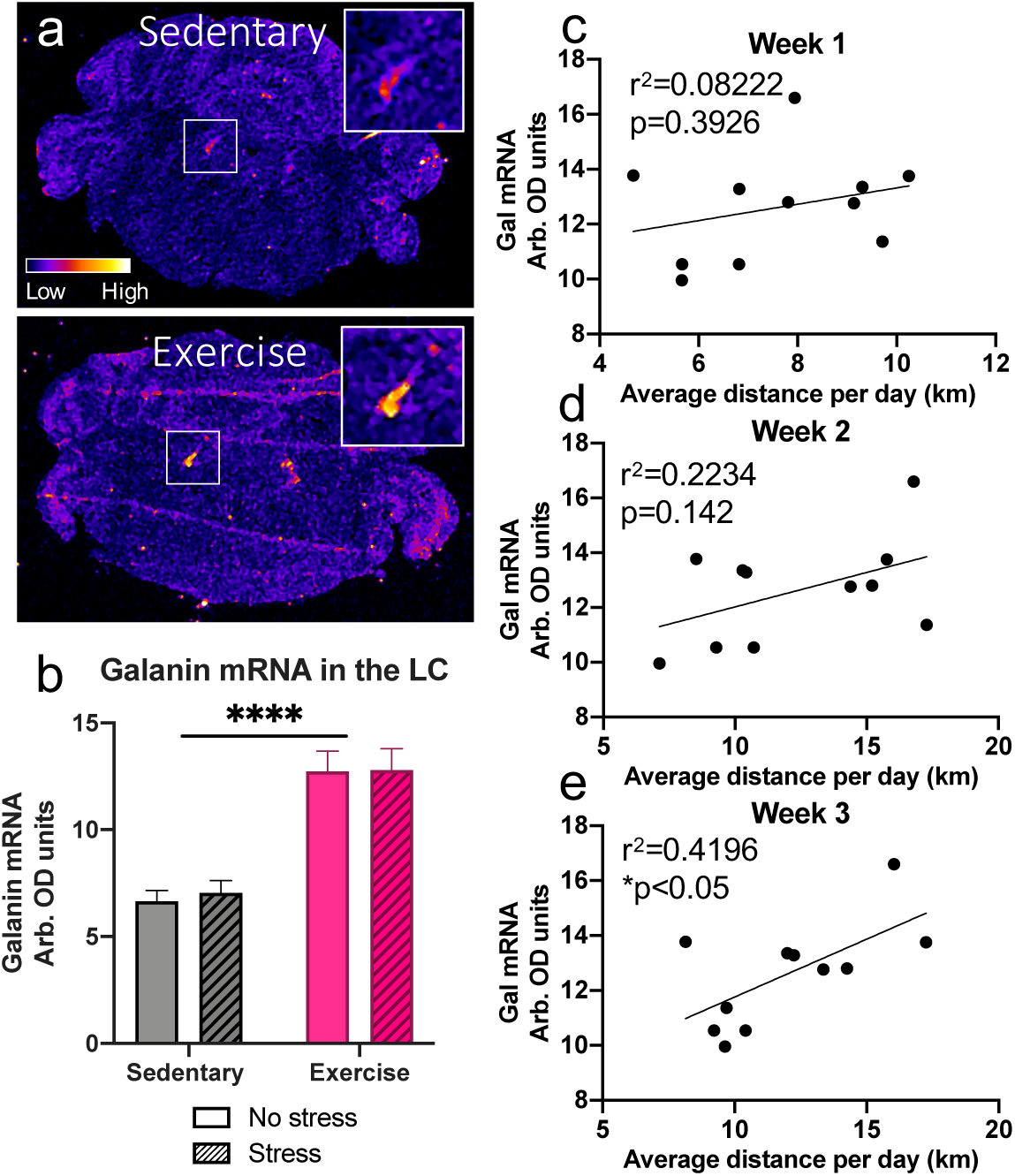
Exercise increases galanin mRNA in the LC of mice. Galanin mRNA levels in the LC were measured via *in situ* hybridization in wild-type C57Bl6/J mice following 3 weeks of *ad libitum* access to running wheels in their home cage (“Exercise”) or no running wheels (“Sedentary”), foot shock (“Stress”) or no foot shock (“No Stress”), and behavioral testing. (a) Representative images of galanin *in situ* hybridization. (b) Quantitative densitometry analysis revealed that exercise mice showed significantly elevated galanin mRNA in the LC compared to sedentary mice, with no effect of foot shock stress exposure. (c) Correlation analysis showed that the average distance ran per day during the third week for each mouse showed a significant positive correlation with the level of galanin mRNA expression in the LC. The average distance ran per day during the first (d) and second (e) weeks did not correlated with the level of galanin mRNA expression in the LC. *n* = 7-8 mice per group. Error bars show SEM. ****p<0.0001.

### Gal OX mice show normal baseline line behavior

Gal OX mice are reported to have normal performance in several canonical tests for anxiety-like behavior, including the elevated plus maze and open field test, but are resistant to yohimbine-induced anxiety-like behavior in a light-dark exploration task (Holmes et al., 2002). To confirm and expand on previous baseline behavioral studies conducted with the Gal OX mice, we conducted a battery of behavioral tasks. We observed no differences between Gal OX and littermate WT control mice in canonical tests of anxiety-like behavior (elevated plus maze, t_17_=0.1641, p=0.8717; zero maze, t_17_=0.6739, p=0.5095; open field, t_17_=0.3211, p=0.7521) or depressive-like behavior (tail suspension test, t_17_=1.578, p=0.1331; forced swim test, t_17_=0.4564, p=0.6539) (**Fig. 4a-e**). They showed no difference from WT littermates in compulsive behaviors during the nestlet shredding task (t_7_=0.07861, p=0.9395) or marble burying task (t_17_=0.8051, p=0.4319) (**Fig. 4f-g**). Additionally, Gal OX behavior was comparable to WT in the novelty-suppressed feeding task (t_17_=1.355, p=0.1933) (**Fig. 4h**). Gal OX mice exhibited normal cognitive responses in contextual (F_1,17_=0.2773, p=0.6053) and cued fear conditioning (F_1,17_=0.5667, p=0.4619) (**Fig. 4i-k**). The locomotor activity of Gal OX mice was normal (F_1,17_=0.6486, p=0.4317). (**Fig. 4l**). In the shock probe defensive burying task, Gal OX mice were normal in most behaviors, including digging (t_12_=0.7124, p=0.4899), freezing (t_12_=0.7015, p=0.4964), rearing, (t_12_=0.4402, p=0.6676), and number of probe touches (t_12_=0.6455, p=0.5308) (**Fig. 5a-d**). The only exception was that Gal OX mice spent significantly less time grooming during the task than their WT counterparts (t_12_=3.152, p=0.0084) (**Fig. 5e**).

**Figure 4.**
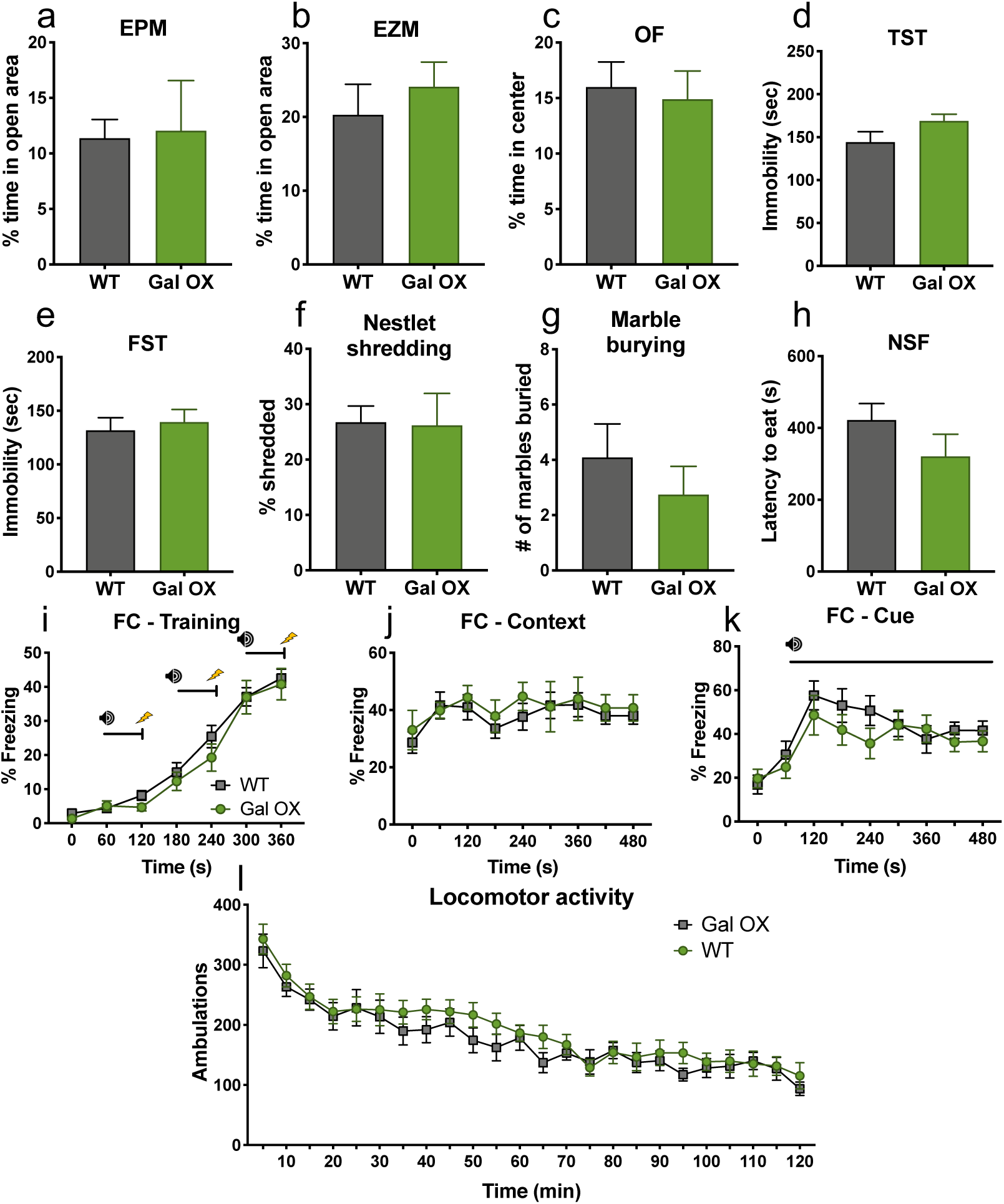
Gal OX mice display normal behavior at baseline. A battery of behavioral tests was conducted on Gal OX mice compared to their WT littermates. (a-b) Open field (OF), (c) marble burying, (d) elevated zero maze (EZM), (e) elevated plus maze (EPM), (f) nestlet shredding, (g) novelty-suppressed feeding (NSF), (h) forced swim test (FST), (i) tail suspension test (TST), (j-l) fear conditioning training, (m) locomotor activity. Gal OX mice were normal in all measures. *n* = 8-11 mice per group. Error bars show SEM.

**Figure 5.**
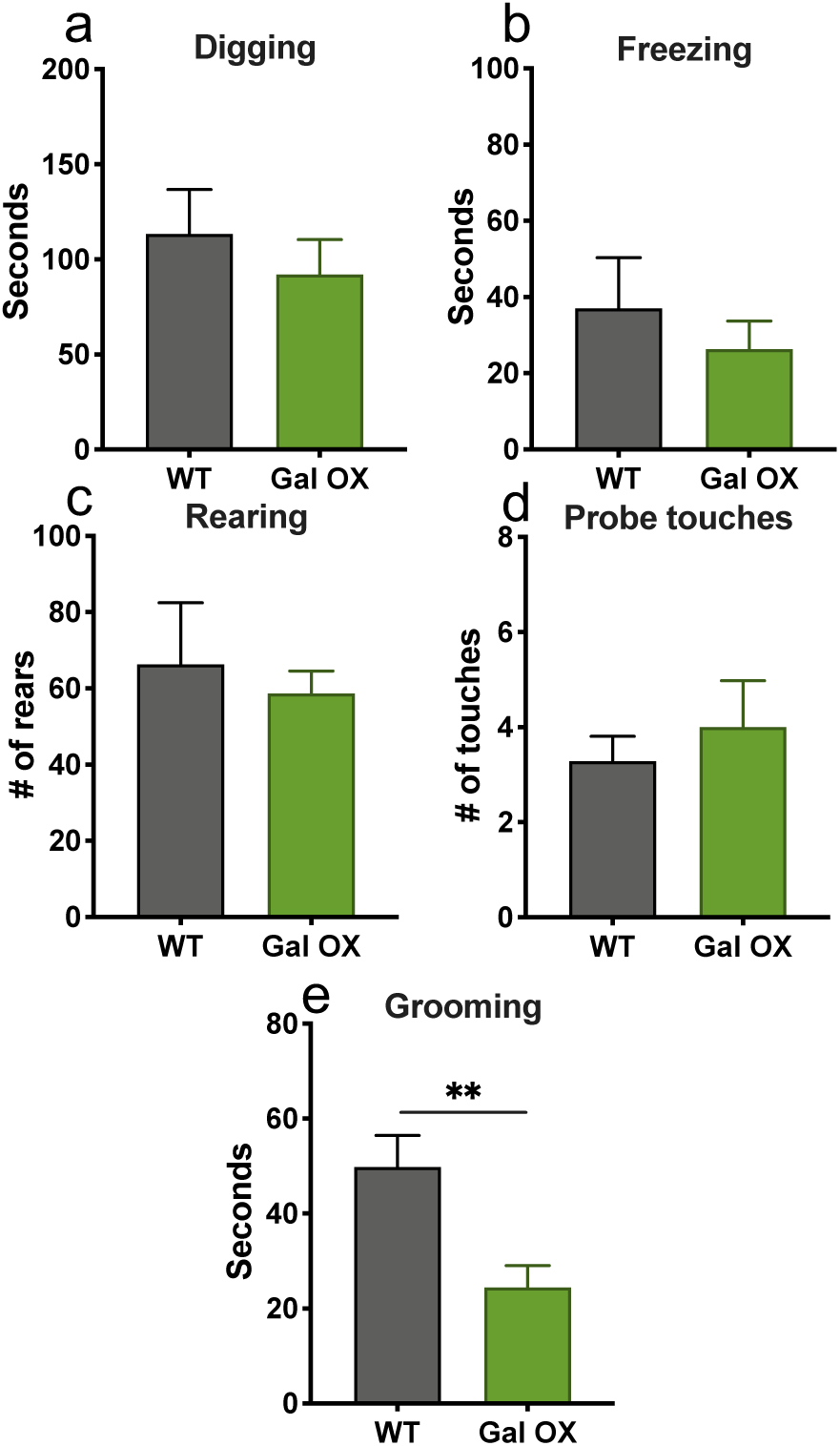
Gal OX mice display decreased grooming in the shock probe defensive burying task. Gal OX mice were normal in all measures in the shock probe defensive burying assay, including time spent digging (a), freezing (b), number of rears (c), and probe touches (d). However, they spent significantly more time grooming than WT controls (e). *n* = 7 mice per group. Error bars show SEM. **p<0.01.

### Gal OX mice are resistant to the behavioral effects of foot shock stress

Although chronic wheel running increased galanin expression in the LC, exercise has pleiotropic effects on brain neurochemistry. To determine whether we could recapitulate the exercise-induced stress resilience effect with chronically elevated noradrenergic galanin expression alone, we tested the response of Gal OX mice to the same foot shock stressor used in the exercise paradigm (**Fig. 6a**). We simplified the paradigm to include only the EZM because the effects of stress and exercise were particularly robust in this task and were representative of the test battery.

**Figure 6.**
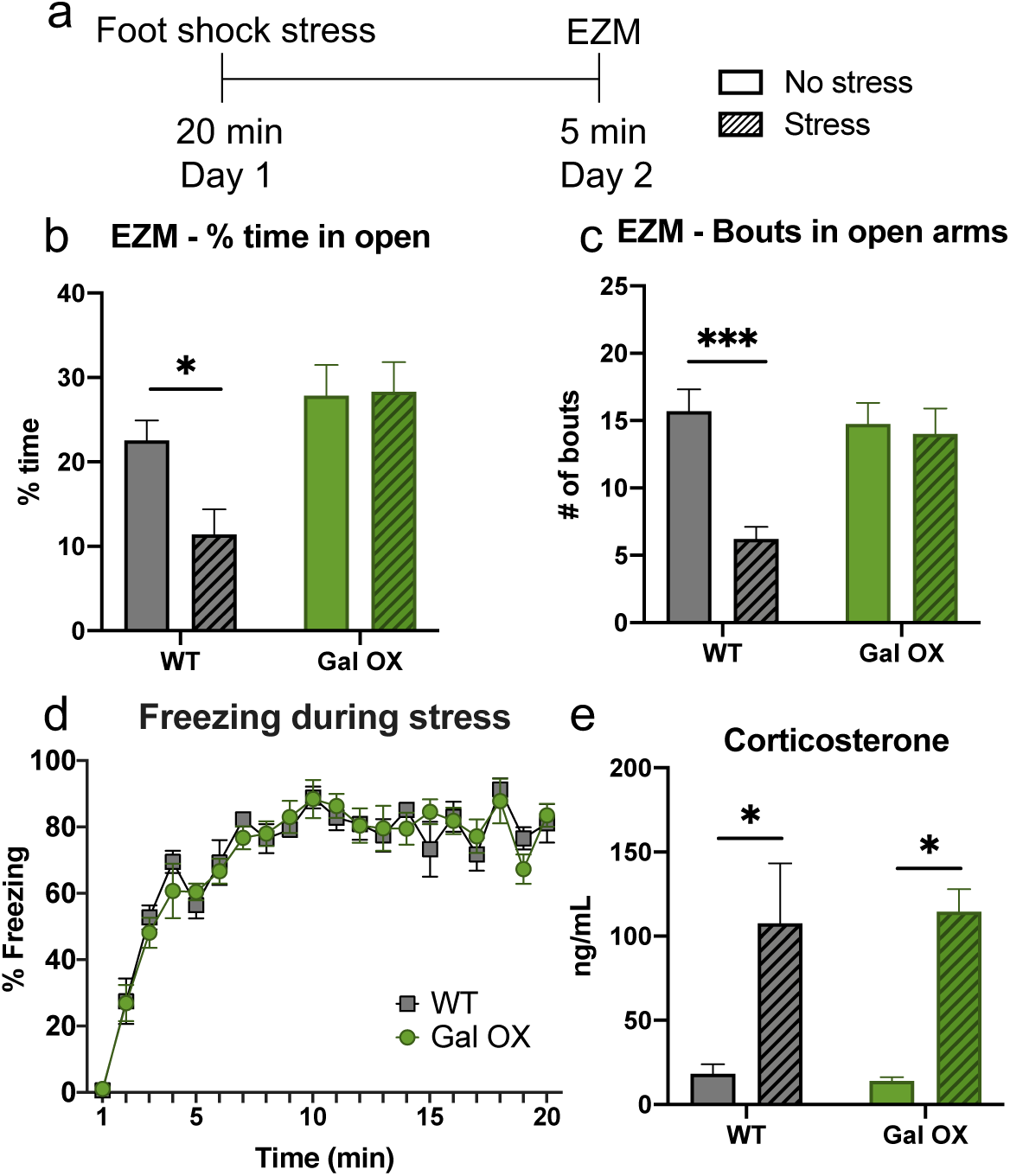
Gal OX mice are resistant to the anxiogenic effects of foot shock stress. Gal OX mice and wild-type littermate controls (WT) received 20 min of foot shock (“Stress”) or no foot shock (“No Stress”) and were tested in the elevated zero maze (EZM) 24 h later. A separate group of mice received foot shock or no foot shock, and blood was collected immediately afterwards for CORT measurements. (a) Foot shock stress paradigm timeline. (b) WT and Gal OX mice showed similar increases in plasma CORT immediately following the foot shock stress. (c) WT and Gal OX mice showed similar freezing behavior during the foot shock stress. WT, but not Gal OX mice, showed a significant decrease in (d) time spent in the open arms and (e) number of open arm bouts in the EZM 24 h after foot shock stress. *n* = 5-7 mice per group for CORT analysis. *n* = 8-10 mice per group for behavior. Error bars show SEM. *p<0.05, ***p<0.001.

When tested in the EZM 24 h after foot shock, there was a main effect of genotype on the amount of time spent in the open arms (F_1,32_=12.25, p=0.0014). WT mice spent significantly less time in the open arms of the EZM 24 h after stressor exposure compared to non-stressed WT mice (p=0.0362). However, Gal OX mice did not show this behavioral change after stress (p=0.9938) (**Fig. 6b**). The same pattern was observed when measuring the number of open bouts in the EZM (F_1,32_=4.659, p=0.0383), where WT mice had significantly fewer open bouts after stress (p=0.0003), but Gal OX mice were unchanged after stress (p=0.9342) (**Fig. 6c**).

We conducted two control experiments to verify that galanin overexpression did not change the physiological responses to foot shock. First, we measured the amount of freezing during the administration of foot shocks and found no differences between WT and Gal OX mice (F_1,10_=0.01299, p=0.9115) (**Fig. 6d**). Next, we measured foot shock-induced increases in plasma corticosterone (CORT) levels and found a significant effect of stress (F_1,23_=17.12, p=0.0004), but no effect of genotype (F_1,23_=0.004147, p=0.9492), indicating that Gal OX mice have a normal CORT response to the foot shock stress itself and the neuroendocrine stress axis is functional in these mice (**Fig. 6e**). These results demonstrate that while Gal OX mice are initially affected by foot shock in the same way as their WT littermates, they are resistant to stress-induced behavioral changes 24 h later.

### Gal OX mice are resistant to the anxiogenic effects of acute optogenetic LC activation

Foot shock stress engages many brain regions in addition to the LC. To more selectively target activation of the noradrenergic system, we used optogenetics to stimulate the LC immediately before EZM testing (**Fig. 7a**). The stimulation and behavioral protocols used here were based on a previously published paradigm of optogenetic LC activation-induced anxiety-like behavior (McCall et al., 2015). Mice received intra-LC infusion of virus expressing either ChR2-mCherry or mCherry alone. Viral expression and optic ferrule placement were confirmed in all animals by histology (**Fig. 7b**), and successful activation of the LC was confirmed via an increase in c-Fos expression (t_8_=2.798, p=0.0233) (**Fig. 7c**). No differences in c-Fos expression after optogenetic stimulation were observed between genotypes. There were significant effects of both genotype (F_1,32_=13.18, p=0.001) and virus (F_1,32_=8.569, p=0.0062) on the amount of time mice spent in the open arms of the zero maze, with post hoc testing showing that WT ChR2 mice spent significantly less time in the open arms compared to WT mCherry mice (p=0.0234), while there was no difference between Gal OX mice that received the ChR2 and the mCherry virus (p=0.2809) (**Fig. 7d**). A similar pattern was seen for the number of open bouts during the EZM test. There were significant main effects of both genotype (F_1,32_=9.64, p=0.004) and virus (F_1,32_=6.463, p=0.0161), with significantly fewer open bouts for WT ChR2 mice compared to WT mCherry mice (p=0.033) and no difference between Gal OX ChR2 or mCherry control mice (p=0.5006) (**Fig. 7e**). These results demonstrate that the Gal OX mice are resistant to LC stimulation-induced anxiety-like behavior.

**Figure 7.**
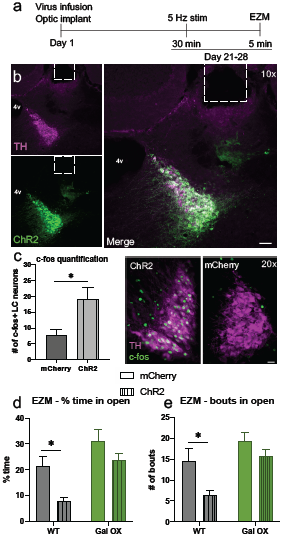
Gal OX mice are resistant to the anxiogenic effects of optogenetic LC activation. Gal OX and wild-type littermate controls (WT) were given 30 min of optogenetic LC stimulation and then tested in the elevated zero maze (EZM). The next day, they were given 15 min of optogenetic LC stimulation, and brains were collected for c-fos immunohistochemistry 90 min later. (a) Optogenetic surgery and behavior timeline. (b) Representative images of ChR2 viral expression (green) and tyrosine hydroxylase (TH, magenta) overlap in the LC and optic fiber ferrule placement (scale bar, 50 µm; 4v, 4^th^ ventricle). (c) Optogenetic stimulation of the LC leads to increased c-fos expression in LC neurons with the ChR2 virus, but not in mice with the control mCherry virus. No differences in c-fos expression between genotypes were seen so data shown are collapsed across genotype (TH, magenta; c-fos, green; scale bar, 20 µm). Optogenetic stimulation causes WT-ChR2 mice to (d) spend less time in the open arms and (e) have fewer open arm bouts in the EZM compared to WT-mCherry mice but does not impact the behavior of Gal OX mice. *n* = 8-10 mice per group. Error bars show SEM. *p<0.05.

## DISCUSSION

Although many studies have suggested a role for LC-derived galanin in stress resilience, most examined a single species (rat) and lacked direct evidence for the noradrenergic system as the functional source of the neuropeptide. To our knowledge, the present results are the first to demonstrate that (1) chronic exercise simultaneously increases both stress resilience and galanin expression in the LC of mice, and (2) increasing galanin in noradrenergic neurons ameliorates anxiogenic-like responses to stress and optogenetic LC stimulation.

We found that the level of galanin expression correlated with running distance in the third week, indicating that the relationship between physical activity and galanin abundance increases in a “dose-dependent” manner, as has been shown in rats (Holmes et al., 2006; Sciolino et al., 2012). For the most part, exercise had no effect on baseline behavior, but ameliorated the anxiogenic effects of foot shock stress, similar to what we have reported in rats (Sciolino et al., 2015). However, we did observe that wheel running increased exploratory drive in behavioral paradigms that were not modulated by stress, such as rearing and probe touches in the SPDB. The results from our chronic voluntary wheel running experiment are consistent with other previous studies that showed enhanced stress resilience after chronic increases in physical activity in rodents (Sciolino et al., 2012; Kingston et al., 2018; Mul et al., 2018; Tanner et al., 2019). Female mice in our study consistently ran farther than male mice, as has been reported previously (De Bono et al., 2006; Bartling et al., 2017), but no other sex differences were seen.

Wheel running has wide-ranging effects on a variety of molecular and structural changes in the brain that may ameliorate the negative affective states induced by stress, including hippocampal neurogenesis and brain-derived neurotrophic factor (Li et al., 2008; Liu and Nusslock, 2018), so it is possible that the alterations in the galanin system as a result of exercise work in concert with other systems in the brain to lead to an increase in stress resilience. To determine whether increasing galanin in noradrenergic neurons, alone, could recapitulate the anxiolytic effects of exercise, we turned to Gal OX mice. Similar to chronic exercise, Gal OX mice showed minimal behavioral differences from their WT littermates at baseline, consistent with much of the previous research suggesting that galaninergic effects are primarily evoked under conditions of stress that elicit high levels of neuronal activity. Galanin, like other neuropeptides, is stored in large dense core vesicles that require high frequency neuronal firing, such as those evoked in LC neurons by stress, to induce release (Lang et al., 2015). Gal OX mice did display decreased grooming in the SPDB assay. Because stress can elicit increased grooming behaviors in mice (Kalueff and Tuohimaa, 2004), decreased grooming could be indicative of increased resilience in the face of a stressful environment like the shock-probe task. Gal OX mice had a typical CORT response to foot shock, indicating that their neuroendocrine stress response was functioning normally, and customary freezing behavior during the stress itself. However, the decrease in open arm time and number of bouts in the EZM evident 24 h after foot shock observed in WT animals were completely abolished in Gal OX mice, suggesting that noradrenergic galanin promotes stress resilience.

While foot shock is a powerful activator of LC neurons, this stressor engages many other nodes of the stress response circuitry (Kim et al., 2016; Lecca et al., 2016). To isolate LC-induced anxiogenesis, we used LC-specific optogenetic stimulation immediately before testing in the EZM. The optogenetic parameters we employed were based on electrophysiological recordings of LC activity in response to stress and previously reported to induce anxiety-like behavior in the EZM. Importantly, the effect of optogenetic stimulation was blocked with β-adrenergic receptor antagonism, demonstrating that increased noradrenergic transmission was responsible for the anxiety-like behavior (McCall et al., 2015). Similar to the foot shock response, Gal OX mice did not display the anxiety-like behavior shown by their WT littermates. The fact that the Gal OX mice are resistant to this behavioral change suggests that increased noradrenergic galanin can dampen the stress-related behavioral effects of increased LC activity.

We consider two main mechanisms by which chronically elevated galanin in the LC leads to increased stress resilience: (1) acute neuromodulatory actions of excess galanin being released during the stressor and acting somatodendritically to inhibit the LC itself or downstream regions involved in mediating stress-induced behaviors, or (2) long-term neurotrophic changes in downstream brain regions mediated by chronically high levels of galanin signaling that confer anxiolysis. All three galanin receptors (GalR1-3) are G protein-coupled receptors that signal through inhibitory G_i/o_ pathways, but GalR2 can also signal through G_q/11_, giving it a more complicated role in downstream signal processing (Lang et al., 2015).

GalR2 is thought to regulate the long-term trophic actions of galanin, whereas GalR1 and GalR3 seem to be more important in the acute inhibitory neuromodulation by galanin (Hobson et al., 2008; Holmes, 2014; Weinshenker and Holmes, 2015). We favor the chronic neurotrophic mechanism to explain our present and previous findings. In our previous study (Sciolino et al., 2015), we found that either chronic exercise or chronic ICV galanin protected against stress-induced anxiogenic behavior and dendritic spine loss in the medial prefrontal cortex (mPFC) of rats. Importantly, chronic but not acute infusion of a galanin antagonist blocked the beneficial effects of exercise on neuroplasticity and behavior, supporting the idea that the stress resilience is caused by chronic elevated galanin leading to long-term neurotrophic adaptation, not acute increases in galanin’s neuromodulatory actions. These data suggest that a similar neurotrophic mechanism likely underlies our present results.

There are several caveats associated with the Gal OX mice used in this study. The first is that the overexpression of galanin is not entirely noradrenergic-specific; they have some ectopic expression of galanin in non-noradrenergic neurons in the piriform and entorhinal cortex (Steiner et al., 2001). Additionally, galanin is not only overexpressed in noradrenergic nuclei that normally express galanin, such as the LC, but all noradrenergic and adrenergic cell groups, leading to high levels of galanin in neurons that do not contain it endogenously. Some of these (e.g. the *Hoxb1* cluster in the medulla and pons outside of the LC (Chen et al., 2019)) have been implicated in stress and anxiety-like responses. Thus, we cannot definitively ascribe the effects we obtained using the Gal OX mice specifically to the LC-derived galanin. Despite this, due to the LC’s well-defined role in regulating the stress response, the broad projections of the LC to other stress-sensitive brain regions, the phenotypic similarities between the Gal OX mice and exercise mice that have elevated galanin exclusively in the LC, and our LC-specific optogenetic results, it seems likely that chronically elevated galanin in the LC contributes to enhanced stress resilience.

Combined with the existing literature, the results from the present study demonstrate that noradrenergic galanin promotes stress resilience across species, and that its enrichment can come from either an environmental (exercise) or genetic (transgenic overexpression) source. Future studies will further dissect how galaninergic transmission from noradrenergic sources underlie its anxiolytic properties. The critical galanin receptor (GalR1-3), the mode of signaling (acute neuromodulatory vs chronic neurotrophic), and the downstream targets in the stress response network (e.g. prefrontal cortex, amygdala) remain to be identified. The galanin system remains a promising target for both pharmacotherapies and non-drug treatments for affective disorders.

## Acknowledgements and author contributions

We thank Jason Schroeder for his technical assistance with the behavioral experiments for this study and Cheryl Strauss for helpful editing of the manuscript. RPT and DW conceived, designed, and supervised the project. Behavioral experiments were performed and analyzed by RPT with assistance from GW. *In situ* hybridization experiments and analysis were performed by RPT with assistance from PVH. Mouse husbandry and genotyping were performed by LCL. RPT and DW wrote the manuscript with input from co-authors.

